# Juvenile mice are susceptible to infarct-induced neurodegeneration that causes delayed cognitive decline in a new model of pediatric ischemic stroke

**DOI:** 10.1101/2025.04.10.648083

**Authors:** Elizabeth W Mayne, Kelly Vanden, Kristy Zera, Marion S Buckwalter

**Affiliations:** Department of Neurology and Neurological Sciences, Stanford University School of Medicine; Division of Child Neurology, Stanford University School of Medicine; Department of Pediatrics, Stanford University School of Medicine; Stanford Stroke Center, Stanford University School of Medicine; Department of Neurosurgery, Stanford University School of Medicine

## Abstract

Over half of pediatric stroke survivors have permanent neurologic and cognitive deficits, and some children develop new or worsening cognitive impairment late after stroke. Their cognitive symptoms are associated with chronic alterations of brain structure in regions distant from the infarct, including in the uninjured hemisphere. However, the mechanisms that cause children’s late cognitive symptoms and disrupted brain growth outside the infarcted tissue remain unknown. We therefore developed and validated a mouse model of pediatric ischemic stroke that causes infarct-induced, delayed cognitive decline accompanied by chronic innate and adaptive neuroinflammation in the infarct and in uninjured subcortical regions that are connected to the infarct.

**Methods:** Male and female C57BL/6J mice were randomized to stroke or sham surgery at p28 to model stroke in late childhood and compared to adult 6-month-old mice. We used permanent distal MCAO followed by 60 minutes of hypoxia, an established model for focal ischemia that generates a purely cortical stroke. Mice underwent behavioral testing at 1- and 7-weeks post stroke Barnes Maze protocol adapted for reversal learning. We performed immunostaining to quantify stroke size, atrophy, and innate and adaptive neuroinflammation at 3 days and 7 weeks after surgery.

**Results:** At 7 weeks post-stroke, juvenile stroke mice performed significantly worse on reversal learning on Barnes Maze testing than sham mice (n = 7 stroke, 7 sham, p = 0.0265 2-way ANOVA with repeated measures). At 3 days after stroke, juvenile mice had greater innate immune cell activation at subcortical sites of secondary neurodegeneration; however, by 7 weeks after surgery, this relationship was reversed, and adults had greater chronic innate immune cell activation at these uninjured subcortical sites (corpus callosum, corticospinal tract, thalamus). Like adult mice, pediatric mice exhibited chronic innate and adaptive neuroimmunity in the infarct scar; the inflammation in the stroke scar did not differ by age.

**Conclusions:** In a model of pediatric ischemic stroke, juvenile mice develop a delayed cognitive deficit that is associated with chronic innate and adaptive neuroinflammation in uninjured subcortical structures that undergo secondary neurodegeneration. Our results suggest that infarct-induced neurodegeneration occurs after stroke in juvenile mice and that juvenile and adult mice have divergent trajectories in the innate immune response to stroke.

## INTRODUCTION

The incidence of pediatric stroke is rising worldwide,^1–3^ and up to 70% of survivors have permanent cognitive impairment that is characterized by prominent deficits in executive function, processing speed, and cognitive flexibility.^4–7^ In children, impairment in these cognitive domains typically presents as a learning disability, and stroke doubles the risk of developing learning disorders such as ADHD to 5.7% in survivors of childhood stroke.^8^ Learning and cognitive disabilities profoundly impair not only children’s quality of life, but also their ability to transition to independent adulthood, with profound psychosocial and financial consequences.^9,10^

Moreover, the emergence of new cognitive symptoms and learning impairments in some survivors years after their stroke is a recognized but poorly understood phenomenon.^11^ In one study, 70% of children with ADHD after stroke had declines in their reading scores over a 4 year period, whereas those with ADHD and no stroke had stable or improved reading scores.^12^ The chronic cognitive deficits after pediatric stroke are associated with widespread morphological changes in brain regions distant from the infarct, such as the cortex and white matter of the uninjured hemisphere.^13–15^ Adult survivors may also have atrophy of uninjured brain regions such as the thalamus that undergo secondary neurodegeneration after stroke, a process known as infarct-induced neurodegeneration.^16–18^ However, some evidence suggests that chronic volume loss in uninjured brain regions may be accentuated in children compared to adults.^19,20^ Minimal autopsy tissue exists from pediatric stroke populations, limiting our ability to infer possible cellular and molecular mechanisms of the widespread changes in brain growth that are observed clinically. Moreover, existing animal work on pediatric stroke has typically focused processes occurring within the infarct itself at acute and subacute timepoints. Thus, the cellular and molecular mechanisms that drive infarct-induced neurodegeneration in children, leading to chronic cognitive disability, altered brain growth, and late cognitive decline after pediatric stroke remain poorly understood.

Given the substantial burden imposed by chronic cognitive dysfunction after pediatric stroke and the paucity of knowledge about its underlying mechanisms, we therefore set out to develop a mouse model of delayed-infarct induced cognitive decline after pediatric stroke, and to determine whether this late cognitive decline was associated with chronic degenerative changes outside the infarct. To identify the specific neurodegenerative mechanisms that drive late cognitive decline (rather than immediate, lesion-related deficits), stroke mice should lack immediate cognitive impairment; thus, any cognitive impairment that emerges at chronic time points will be attributable to infarct-induced neurodegeneration. The cognitive domains impaired in the model should also align with those observed in pediatric stroke survivors (e.g., processing speed, executive function, cognitive flexibility), while sparing functions not typically affected after pediatric stroke (e.g., spatial memory). Finally, it should induce neurodegeneration in brain regions that are not acutely injured but which receive or send projections to the infarcted tissue, an essential requirement to interrogate the mechanisms that cause extensive changes in development throughout the entire brain after childhood stroke.^13,15^

## METHODS

### Animals

All mice were male C57BL/6J mice purchased from Jackson Laboratories (Catalog 000664). Juvenile mice were postnatal day 28 and adult mice were aged 6 months at the time of surgery. Mice were group housed up to 5 mice per cage on a 12-hour light/dark cycle, with ad libitum access to food and water. All procedures were approved by the Stanford University Institutional Animal Care and Use Committee.

### Experimental Procedures

#### Distal Middle Cerebral Artery Occlusion Stroke Model (dMCAO stroke)

Distal middle cerebral artery occlusion (dMCAO) stroke was performed as previously described.^21^ In brief, the scalp was shaved and mice were anesthetized using isoflurane at 2.0 L/min. Temperature was maintained at 37ºC throughout the surgery using a homeothermic heating system with a rectal probe to monitor core temperature (Stoelting Catalog #50300). Once the mouse was anesthetized, the scalp was disinfected with chlorhexidine surgical scrub and ophthalmic eye ointment was placed over the eyes. An incision was made over the skull midline and anterior to the right ear. The right temporal muscle was mobilized at the superior border, exposing the underlying skull. A surgical microdrill (Harvard Apparatus catalog 72-6065) was used to make a small craniotomy overlying the distal middle cerebral artery. The dura was incised and cleared off the artery. For stroke surgeries, the artery was then cauterized with a Bovie DEL0 Change-A-Tip Deluxe Low Temperature Cauterizer. All steps were identical for sham surgery, except for cauterization of the artery. The area was then irrigated by sterile saline to wash away any debris or blood. The temporalis was replaced and the surgical wound was closed with surgical glue Meridian Surgi-Lock 2oc Instant Tissue Adhesive. All mice received subcutaneous injections of 0.5 mg/kg Buprenorphine ER (0.5 mg/ml, Wedgewood Veterinary Pharmacy) and cefazolin 25 mg/kg (Cefazolin sodium salt, VWR catalog 89149-888) following surgery.

#### Distal Hypoxic Stroke (DH stroke)

DH stroke was performed identically to dMCAO stroke, then followed by hypoxia.^22^ After wound closure, the animal was allowed to recover for 5 minutes following the end of anesthesia, then placed in a hypoxia chamber (Coy vinyl anaerobic chamber; Coy Laboratory Products) containing 8% oxygen / 92% nitrogen for 60 minutes. Temperature in the hypoxia chamber was maintained at 37 ºC.

### Behavioral Testing

All mice were group housed to avoid stress from social isolation. Testing was performed during the light phase without food or water restriction and recorded with an overhead Logitech C930e 1080P HD Webcam. Barnes maze^23^ and novel object recognition (NOR)^23,24^ tests were performed at 1 and 7 weeks after surgery. For Barnes Maze, we tested two protocols in independent cohorts of mice. In the Barnes maze reversal protocol, mice underwent 3 trials per day for 3 days of training; the release location was moved for each trial, and the location of the escape hole was moved each day to emphasize reversal learning and cognitive flexibility. For the Barnes maze spatial learning paradigm, mice underwent 3 trials per day for 3 days of training; the animal was released in the center of the maze for every trial, and the location of the escape hole remained constant throughout each time point. Automated behavior scoring software (BehaviorCloud, BehaviorCloud LLC) was used to measure time to locate the escape hole, distance traveled, and average speed. NOR was performed as previously described with a fixed 2 hour intertrial interval.^23–25^ Open field testing was performed as previously described,^23^ and BehaviorCloud was used to measure center crossings, time in the center, average speed, and path length.

### Tissue Preparation

All mice were terminally anesthetized with a ketamine/xylazine cocktail, then perfused transcardially with ice cold PBS. After perfusion, brains were drop fixed in 4% PFA in phosphate buffer for 24 hours at 4ºC, followed by impregnation with 30% sucrose. Forty µm thick coronal brain sections were prepared using a freezing microtome (Microm HM430) and stored at -20ºC in a cryoprotective solution containing 30% glycerin, 30% ethylene glycol, and 40% 0.5M sodium phosphate buffer).

### Immunostaining

Immunostaining was performed on PFA-fixed brain sections as previously described.^26^ B lymphocytes were stained with biotinylated anti-mouse B220 (1:1000, BD Biosciences 553085) and T lymphocytes with anti-mouse CD3ε (1:500, BD Biosciences 550277). Activated myeloid cells were stained with anti-mouse CD68 (1:1000, BIO-RAD, MCA1957). Reactive astrocytes were stained with anti-mouse GFAP (1:10,000, Agilent Dako Z0334). Neurons were stained with mouse anti-NeuN (1:500, Sigma-Aldrich MAB377). Secondaries were biotinylated goat-anti-hamster (BA-9100-1.5), goat-anti-rabbit (BA-1000-1.5), and rabbit anti-rat (BA-4001-.5) at 1:500-1:1000 from Vector Laboratories.

### Imaging

All images were acquired on Keyence BZ-X700 series inverted bright field microscope. Quantification of staining in the infarct core and corpus callosum was done in 5 sections per mouse, and quantification of the thalamus and corticospinal tracts was done in 2 sections per mouse. Anatomic regions were located using the Allen Mouse Brain Atlas. Automated thresholding using a custom ImageJ macro was performed to reduce bias. For infarct volume analysis, every 6th 40 µm thick section was stained using NeuN. The infarct core, ipsilateral hemisphere, and contralateral hemispheres were manually traced. Infarct volume was calculated using the Cavalieri method^27^ and reported as % of the ipsilateral hemisphere.

### Statistical analysis

Normality testing was performed using GraphPad Prism 9 and nonparametric tests were used where appropriate. Comparisons of 2 groups at a single timepoint were done using Student’s t-test (parametric) or Mann-Whitney (non-parametric). Comparisons of 3 or more groups were done with 2-way ANOVA, followed by Šidák correction by post-hoc pairwise comparisons. Learning over the training period of Barnes Maze was analyzed using a 1-way ANOVA.

## RESULTS

We used permanent distal middle cerebral artery occlusion (dMCAO stroke) to induce stroke because this model does not cause motor impairment (which could confound behavior testing) and spares the hippocampus (which would produce immediate cognitive impairment that could mask or confound cognitive decline that emerges over time as a consequence of infarct-induced neurodegeneration). The immature brain may have greater density of blood vessels and thus more robust collateral blood flow than the adult brain,^28–30^ so stroke models in juveniles might not produce the same infarct volumes as in adults. Therefore, we began by quantifying infarct volumes in two permanent distal occlusion models that produce delayed cognitive deficits in older mice: distal middle cerebral artery occlusion (dMCAO)^21^ and distal hypoxic stroke (DH stroke),^22^ in which the permanent dMCAO is followed by 60 minutes of hypoxia, resulting in larger and more consistent infarct volumes in adult C57BL/6J mice.

### Juvenile DH stroke induces a reproducible, cortical stroke with low variability, low mortality and comparable size to adult DH stroke

We quantified infarct volume acutely at 72 hours and chronically at 7 weeks after stroke (when cognitive impairment emerges in the adult model of infarct-induced delayed cognitive decline^26^). Juvenile dMCAO stroke produces very small strokes (Figure 1b, n = 7), and in up to 20% of animals, no stroke at all (6/28 animals over 3 experiments), which is undesirable because it would introduce significant variability in behavior results. The addition of 1 hour of hypoxia after stroke or sham surgery increases acute stroke size to a volume comparable to that in adult (6mo) mice that undergo DH stroke (Figure 1b). The hypoxia does not cause neuronal loss in sham-operated animals or outside the infarct in stroke-operated adults.^22^ Over time, DH stroke leads to volume loss in the ipsilateral hemisphere, which is consistent between age groups (Figure 1c). Mortality with DH stroke surgery in juveniles is also low, 15 ± 5%. We thus decided to move forward and test whether juvenile DH stroke produces delayed cognitive deficits.

**Figure 1:**
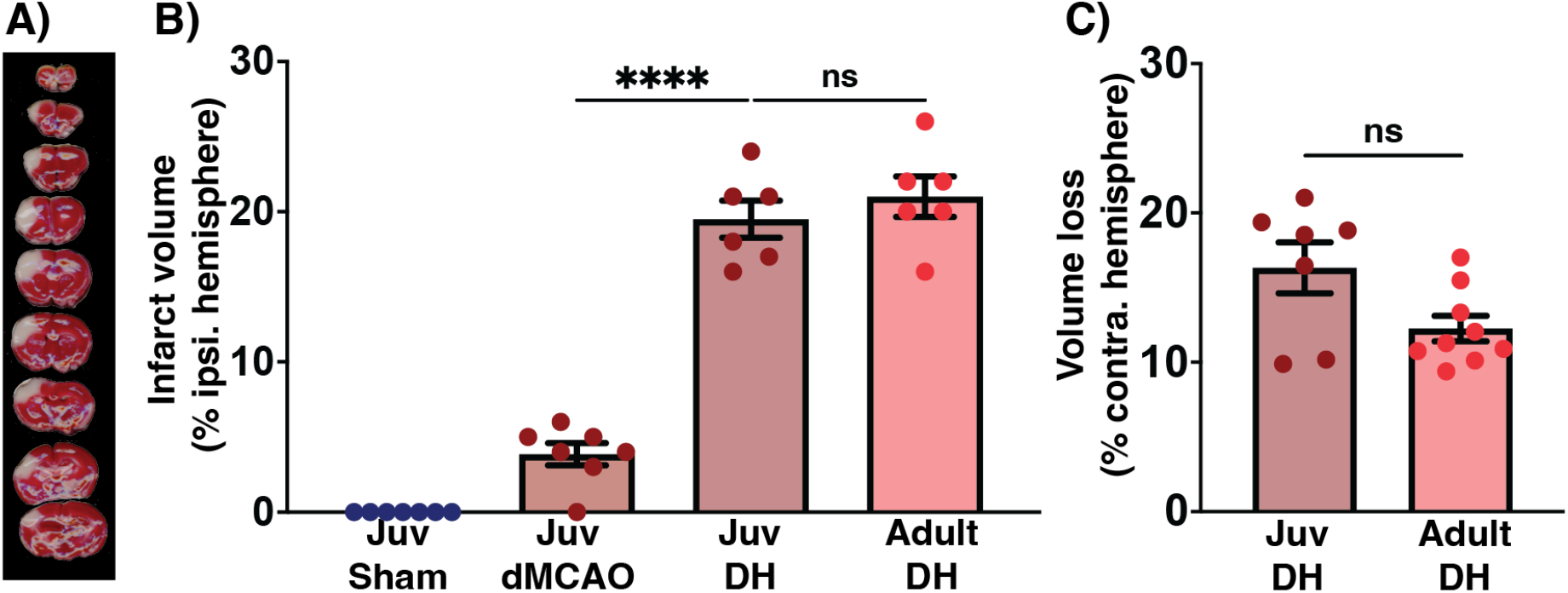
Juvenile DH stroke. A) TTC staining of juvenile DH stroke at 72 hours after surgery. B) Acutely (72 hrs after surgery), juvenile dMCAO stroke produces small infarcts, while juvenile DH stroke produces larger infarcts comparable in volume to adult animals (n=6-7/group, 1-way ANOVA with Šidák correction for multiple comparisons). C) At 7 weeks after surgery, DH stroke induces volume loss in the ipsilateral hemisphere of both juveniles and adults (n=7-9/group, Mann-Whitney). All graphs mean ± s.e.m. * padj < 0.05; ** padj < 0.01; *** padj < 0.001; **** padj < 0.0001.

### Juvenile DH stroke causes a delayed, infarct induced impairment in reversal learning

To evaluate for delayed cognitive decline in juvenile DH stroke, we tested cognition at both an acute time point (1 week after stroke) and a chronic time point (7 weeks after stroke). To interrogate cognitive function after stroke, we used the Barnes maze and novel object recognition tasks. We used two different Barnes maze protocols in independent groups of mice. The first protocol tested testing reversal learning (a correlate of cognitive flexibility,^31^ which is often impaired in pediatric stroke survivors). The second protocol tested spatial memory (which is relatively spared in pediatric stroke survivors).^5^ This allowed us to determine whether the model recapitulates the specific cognitive domains affected after childhood stroke.

In the reversal learning paradigm (Barnes maze-reversal), the mice must locate the escape hole in a new location each day and are released at a different position on the maze for each trial (Figure 2a). One week after surgery, there was no difference between sham-operated and stroke-operated mice in performance on the Barnes maze reversal paradigm. However, by 7 weeks after surgery, the stroke-operated mice developed a reproducible deficit in reversal learning and performed progressively worse throughout the training period (Figure 2a).

**Figure 2:**
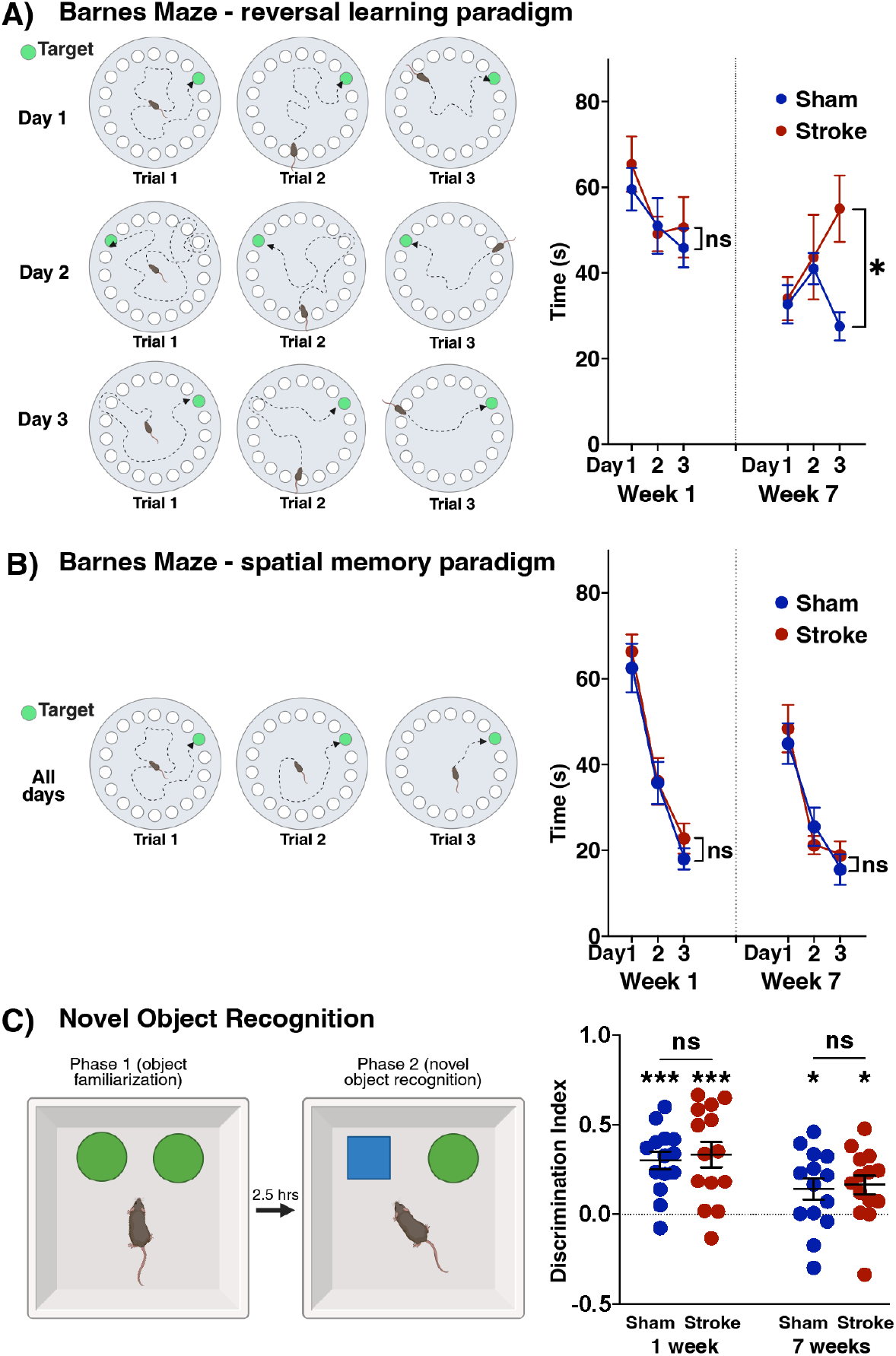
Juvenile DH stroke induces a delayed, selective impairment in reversal learning. A) Left: Schematic of the reversal learning paradigm. The release location changes on each trial, and the location of the escape hole changes each day. Right: quantification of the mean time to locate the escape hole. Juvenile stroke and sham mice perform comparably 1 week after stroke (p = 0.4322, F(1,12) = 0.6, two-way ANOVA with repeated measures, n = 7/group). However, at 7 weeks after stroke, the stroke mice have a deficit in reversal learning and perform significantly worse than sham mice, n = 7/group, p = 0.0428, *F*(1,12) = 5.1, two-way ANOVA with repeated measures. B) Left: Schematic of the spatial memory paradigm. Both the release location and the escape hole location are fixed across all trials and all days of training. Right: Juvenile stroke and sham mice perform comparably at both 1 and 7 weeks after stroke on the spatial memory task. (Week 1 *F*(1, 27) = 0.7983, p = 0.3795); week 7 (*F*(1, 27) = 0.04952, two-way ANOVA with repeated measures, p = 0.8256, n=14 sham, 16 stroke). C) Left: Schematic of novel object recognition task. Right: Juvenile stroke and sham mice both exhibit a preference for the novel object, indicated by DI > 0 at 1 and 7 weeks after stroke, n = 14-15/group, Wilcoxon rank sum test. There is no difference between stroke and sham groups at either time point, n = 14-15/group, Mann-Whitney. * p < 0.05. ** p < 0.001

There was no difference in average speed on open field testing at 1 or 14 weeks after stroke in a separate cohort of mice (n = 14 sham, n = 16 stroke, Supplemental Figure 1). Thus, the deficit in stroke mice on reversal learning at 7 weeks does not reflect fatigue or weakness.

We next tested whether the delayed cognitive impairment in stroke-operated mice was specific for cognitive flexibility. We used a Barnes maze paradigm in which both the release location of the mouse and the position of the escape hole remain fixed throughout all trials at a given time point (Figure 2b). This paradigm (Barnes maze-spatial) evaluates spatial and short-term learning and memory and does not require reversal learning. There was no difference between stroke-operated mice and sham-operated mice at either 1 or 7 weeks after surgery on the Barnes maze spatial paradigm. The novel object recognition task also tests short term memory. Both stroke and sham-operated groups exhibited a preference for the novel object at both time points (median discrimination index significantly different from 0). Congruent with the results on the spatial learning version of the Barnes maze, we also did not observe a difference between stroke- and sham-operated mice on the novel object recognition task (Figure 2c). The discrimination index for all groups was significantly different from 0 (no preference for the novel object) on Mann-Wilcoxon rank testing.

Both the adaptive and innate immune responses to stroke are implicated in post-stroke cognitive decline in adult humans and mice.^26,32–35^ However, it is unknown whether children or juvenile mice have chronic neuroinflammation after stroke, particularly in brain regions outside the infarct. Additionally, the immature immune system in children differs from adults.^36^ Therefore, we next compared the acute (72 hour) and chronic (8 week) inflammatory responses to stroke between juveniles and adults in uninjured subcortical structures (thalamus, corpus callosum, and corticospinal tracts).

### There is greater acute myeloid cell inflammation at sites of secondary neurodegeneration in juvenile stroke than adult stroke

At 72 hours after stroke, there is acute neuroinflammation within the infarct and peri-infarct cortex characterized by presence of reactive astrocytes and activated myeloid cells at the border of the infarct. There was no difference between juveniles and adults in CD68 staining for activated myeloid cells within the infarct or peri infarct cortex (n = 13 juveniles, 4 adults, Mann-Whitney testing, Supplemental Figure 2). Similarly, there was no difference in GFAP staining for reactive astrocytes in the infarct (n = 12 juveniles, 5 adults, Mann-Whitney testing, Supplemental Figure 2).

By 72 hours after stroke, Wallerian degeneration is initiated, and death of connected axons leads to innate immune activation in adult mice.^37^ Interestingly, there was significantly more myeloid cell inflammation as measured by CD68 and Iba1 immunoreactivity in juveniles in regions that are highly connected to the stroke and thus expected to have secondary / Wallerian degeneration after the infarct. This included the corpus callosum, thalamus, and corticospinal tracts (Figure 3a-c, n=6-11/group). Unexpectedly, juveniles had greater CD68 immunoreactivity per µm^2^ in the contralateral corpus callosum, Figure 3a. Similarly, there was greater CD68 reactivity in the juvenile corticospinal tract (Figure 3b) and juvenile thalamus (Figure 3c). Iba1 staining showed a similar increase in staining in juveniles as compared to adults (Supplemental Figure 3). The increased microglial response in connected regions was not mirrored in astrocytic GFAP expression, which was not different (Supplemental Figure 4).

**Figure 3:**
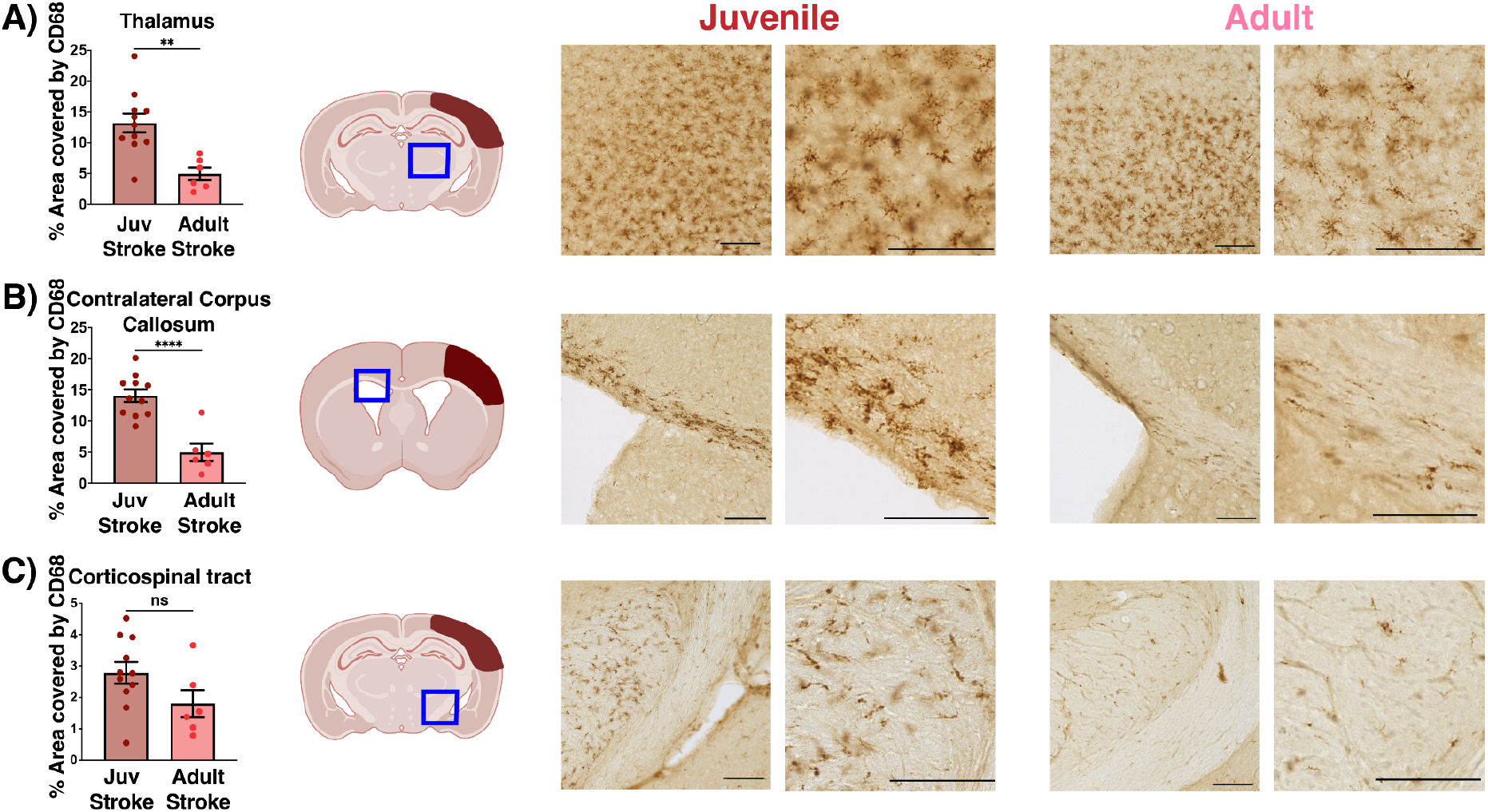
There is greater acute myeloid cell activation at sites of secondary neurodegeneration in juvenile mice at 72 hours after DH stroke. A) Left: quantification of CD68 staining in the ipsilateral thalamus (denoted by blue box in middle panel). Juveniles have significantly greater CD68 immunoreactivity than adults. B) Quantification of CD68 staining in the contralateral corpus callosum (denoted by blue box in middle panel). Juveniles have significantly greater immunoreactivity than adults. C). There is no significant difference between juveniles in adults in CD68 immunoreactivity in the ipsilateral corticospinal tract. Scale bar 100 µm. Statistics: Mann-Whitney test, n = 11 juveniles; 6 adults. Quantification denotes mean ± S.E.M. Mann-Whitney test. * p < 0.05; ** p < 0.01; *** p < 0.001; **** p < 0.0001.

### Juvenile DH stroke causes attenuated chronic myeloid cell neuroinflammation in regions of secondary neurodegeneration compared to adult stroke

Given the unexpected findings that stroke elicited greater chronic myeloid cell activation in the connected brain regions of juvenile mice than adult mice, we next asked whether this resulted in differences in chronic myeloid inflammation between juveniles and adults. We sacrificed juvenile and adult mice at 7 weeks after surgery and quantified myeloid cells in the same regions as Figure 3.

Both juvenile and adult stroke animals had greater myeloid cell activation in the corpus callosum, corticospinal tract, and thalamus than sham animals (Figure 4a-c, n=6-7/group). When comparing juvenile stroke mice to adult stroke mice, the juvenile stroke mice had significantly less CD68 staining in the thalamus (Figure 4a) and corticospinal tract (Figure 4c), and a non-significant trend toward less CD68 staining in the corpus callosum (Figure 4b).

**Figure 4:**
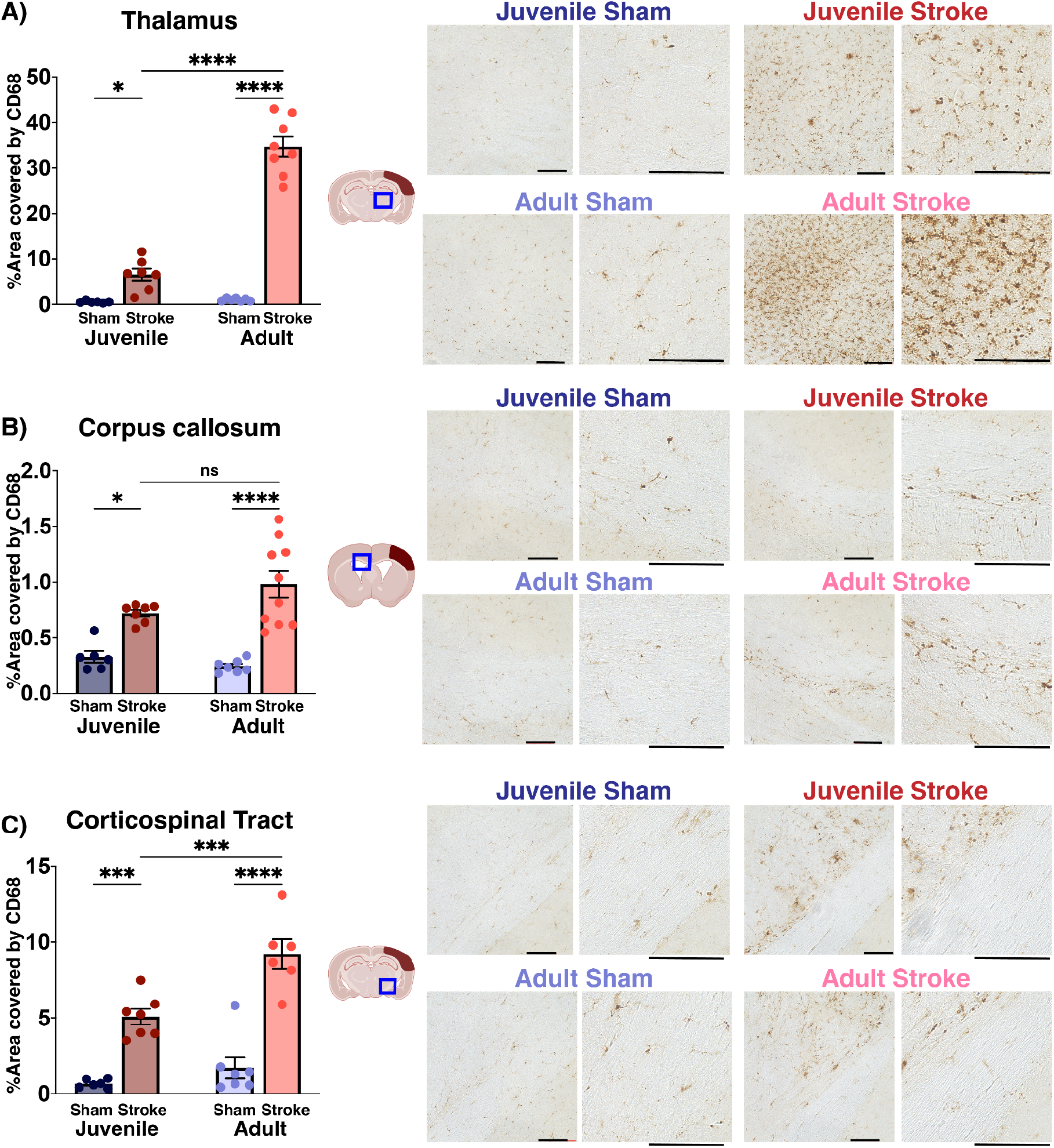
Both juvenile and adult mice have chronic myeloid cell immunoreactivity at sites of secondary neurodegeneration, but it is attenuated in juveniles. A) There was significantly less CD68 immunoreactivity in the ipsilateral thalamus of juvenile stroke mice compared to adult stroke mice. B) There was a nonsignificant trend toward diminished CD68 immunoreactivity in the contralateral corpus callosum of juveniles compared to adults. C) There was significantly less CD68 immunoreactivity in the ipsilateral corticospinal tract of juvenile mice compared to stroke mice. Scale bar 100 µm. Statistics: One-way ANOVA with Šidák correction for multiple comparisons, n = 6-7/group. * p < 0.05; ** p < 0.01; *** p < 0.001; **** p < 0.0001. Error bars denote mean ± S.E.M.

### Juvenile DH stroke causes comparable chronic lymphocytic neuroinflammation in the infarct

Because chronic B lymphocyte neuroinflammation has previously been implicated in infarct-induced cognitive decline,^26,32,38^ we next determined whether juvenile stroke causes chronic lymphocytic neuroinflammation. Both juvenile dMCAO and DH stroke cause chronic B and T lymphocyte neuroinflammation in the infarct scar that persists for at least 7 weeks after stroke. There was not a significant difference in chronic B lymphocyte or T lymphocyte neuroinflammation between juvenile stroke mice and adult stroke mice (Figure 5a-b).

**Figure 5:**
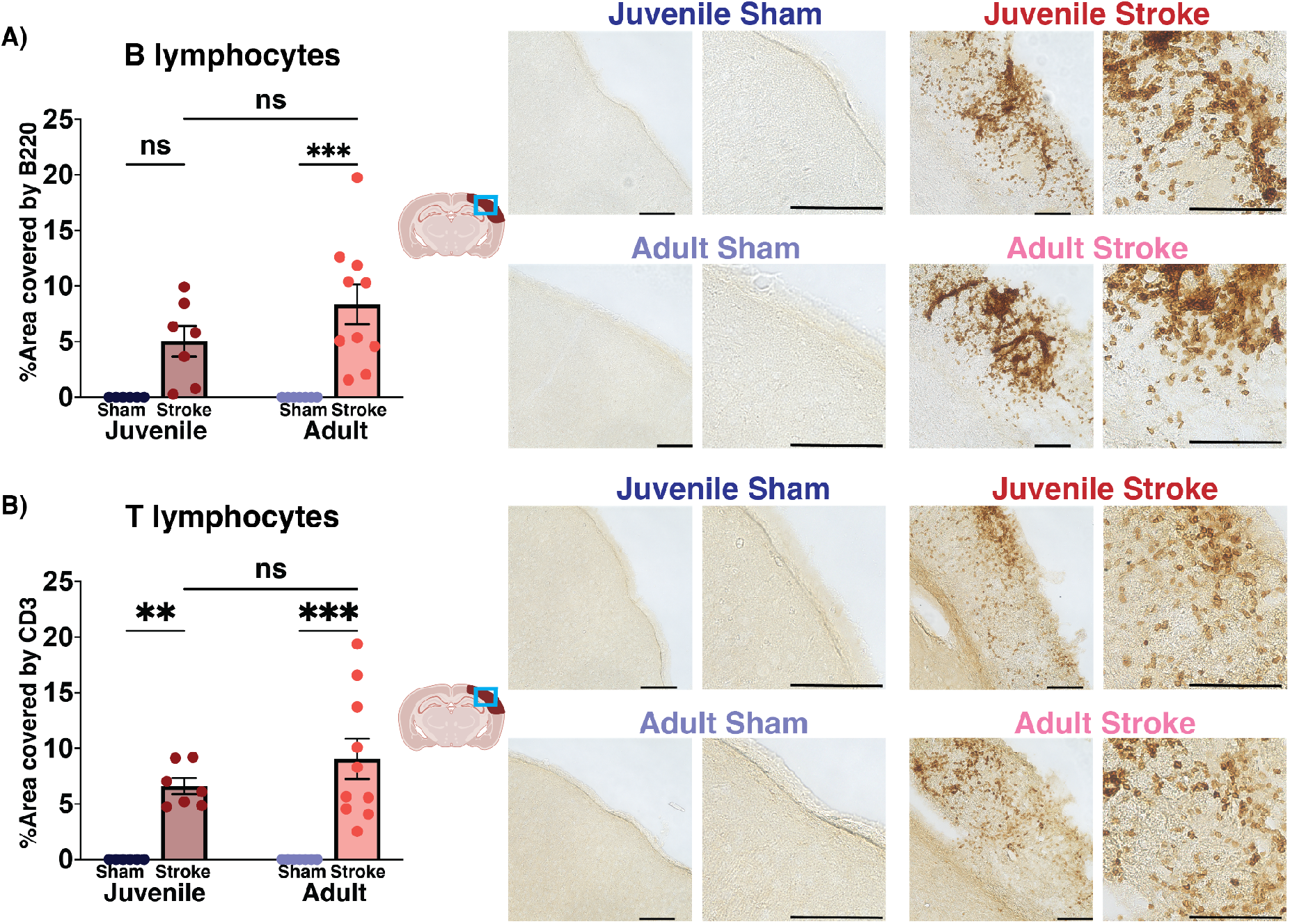
Juvenile and adult mice have comparable chronic lymphocytic neuroinflammation after stroke. A) Quantification of B220 immunostaining for B lymphocytes shows that both juveniles and adults have chronic B lymphocyte neuroinflammation in the infarct scar at 8 weeks after stroke. B) CD3 immunostaining for T lymphocytes shows chronic T lymphocyte neuroinflammation in the infarct at 8 weeks after stroke in juvenile and adult mice. Scale bar 100 µm. Statistics: One-way ANOVA with Šidák correction for multiple comparisons, n = 6-7/group. * p < 0.05; ** p < 0.01; *** p < 0.001; **** p < 0.0001. Error bars denote mean ± S.E.M.

## DISCUSSION

Here, we present a novel model for juvenile, infarct-induced cognitive decline accompanied by acute and chronic neuroinflammation both within the infarct and in uninjured subcortical regions that send or receive projections from the infarct and undergo secondary neurodegeneration. Intriguingly, while juveniles and adults have comparable acute myeloid and astroglial inflammation within the infarct, juveniles have markedly greater myeloid cell activation in uninjured subcortical structures including the thalamus, corpus callosum, and internal capsule. This relationship reverses at chronic time points, when adults have greater myeloid neuroinflammation in subcortical structures. However, there is no difference in lymphocytic neuroinflammation between juveniles and adults.

The finding that the developing brain is susceptible to infarct-induced cognitive decline is a major shift from the historical conceptualization of pediatric stroke as a static brain injury. It also suggests that cognitive testing in juvenile stroke models with earlier time points may miss this delayed cognitive decline,^39^ highlighting the need for models of chronic cognitive function in a disease where the majority of survivors are expected to live for decades after stroke. Crucially, our model recapitulates deficits in the cognitive domains that are typically affected after stroke in children -namely, cognitive flexibility and processing speed.^4,5,11,40^ Notably, most commonly used rodent behavioral tasks prioritize hippocampal-dependent processes such as spatial or short term memory and are less sensitive for deficits in executive function and processing speed.

Infarct-induced autoimmunity is strongly implicated in driving delayed cognitive decline in adults. We show that juvenile stroke induces chronic innate and adaptive neuroinflammation, both within the infarct and in connected white matter tracts and deep gray nuclei outside the infarct, which has not previously been reported in juvenile models. This finding is similar to that reported in adults, where it has been attributed to Wallerian degeneration.^26,41,42^ In human adults and children, MRI imaging also detects changes in connected regions consistent with Wallerian degeneration,^16,18–20^ but remote inflammation has not been demonstrated previously in models of juvenile stroke at either acute or chronic timepoints, and our results suggest a potential mechanism by which focal injury can drive the wider disruption of both gray and white matter structures that have been reported clinically.^13,15,19,20^

Finally, we observe chronic lymphocytic inflammation in the infarct that is similar between juveniles and adults, which is an important finding because B lymphocyte-mediated neuroinflammation mediates delayed, infarct-induced neurodegeneration in adult mice, and depleting B lymphocytes prevents cognitive decline. This identifies a possible mechanism with potential for therapeutic intervention, as B lymphocyte depleting therapies are already in widespread use in pediatric neuroimmune diseases.^43,44^

While our study identified important similarities between the juvenile and adult brain after stroke, it also highlighted crucial differences. First, while juvenile mice in our study also develop infarct-induced cognitive decline, the cognitive domains differed from those reported in adults. Adult mice develop infarct-induced, progressive deficits in hippocampal long term potentiation in parallel with emerging impairment on hippocampal dependent tasks such as novel object recognition,^26^ In contrast, the juvenile mice in our study developed a reproducible deficit selectively in reversal learning, a correlate of cognitive flexibility. This matches the cognitive deficits that are reported in survivors of childhood stroke, who often have profound impairment in executive function and processing speed, while spatial and short term memory are relatively spared.^4,5^ This highlights the validity of our model, and also suggests that there are likely important differences in the mechanisms of infarct-induced cognitive decline between the developing and adult brain.

We also observed striking age-related differences in the trajectory of innate immunity at sites of secondary neurodegeneration after stroke. While acute myeloid and astroglial inflammation within the infarct and peri-infarct cortex was similar between juveniles and adults, the acute myeloid cell inflammation outside the infarct was more pronounced in juveniles compared to adults. Intriguingly, this relationship reversed at chronic time points, when adult mice showed greater persistent innate neuroimmunity in these sites of secondary neurodegeneration. Innate neuroinflammatory responses at *uninjured* sites of secondary neurodegeneration have not been previously examined at either acute or chronic time points in juvenile models of stroke, nor have those trajectories been compared to adult stroke.

The age-related differences we observed in the trajectory of activated myeloid cells sites of secondary neurodegeneration highlight that there are likely important differences in the innate immune to stroke between juveniles and adults. Given the heterogeneity of microglial functions across age and disease states, it remains unknown whether the increased CD68 staining we observed in connected white matter tracts in juveniles as compared to adults is maladaptive or protective. Future studies with single-cell resolution using RNA sequencing and/or flow cytometry will provide additional insight into whether there are also important functional or phenotypic differences between the myeloid cells that participate in the chronic innate response to stroke in the juvenile and adult brain. Because transcriptionally distinct microglial subpopulations likely have different functions, this could be an important driver of divergent outcomes after juvenile and adult stroke, and this is an important direction for future work.

Cognitive impairment is common after pediatric stroke, affecting over half of survivors. Stroke roughly doubles the risk of developing ADHD and other learning disabilities, which has long-lasting implications for children’s health, quality of life, and transition to independence as adults.^45^ As in adult post-stroke cognitive impairment, some children appear to be more vulnerable,^46^ and a subset of children may worsen over time.^12^ However, the cellular mechanisms that cause children’s cognitive deficits after stroke remain unknown. Because of the challenges in conducting clinical research into children’s long-term cognitive outcomes and the lack of autopsy tissue, identifying these mechanisms requires development of translational models that complement our clinical studies. Here, for the first time, we report that focal stroke induces chronic, widespread neuroinflammation in the juvenile brain, and that this is associated with infarct-induced cognitive decline. This model will support investigation into the mechanisms that drive chronic cognitive dysfunction after pediatric stroke. Identifying therapeutic targets in the subacute and chronic period after pediatric stroke is of paramount importance, since very few children are eligible for treatment with the hyperacute revascularization therapies (tPA/TNK, EVT) that have revolutionized outcomes in adult stroke.

## Non-standard Abbreviations and Acronyms

dMCAO stroke: distal middle cerebral artery occlusion stroke
DH stroke: distal hypoxic stroke
GFAP: glial fibrillary astrocytic protein
NOR: novel object recognition test

