## Supplemental Figures for "Juvenile mice are susceptible to infarct-induced neurodegeneration that causes delayed cognitive decline in a new model of pediatric ischemic stroke"

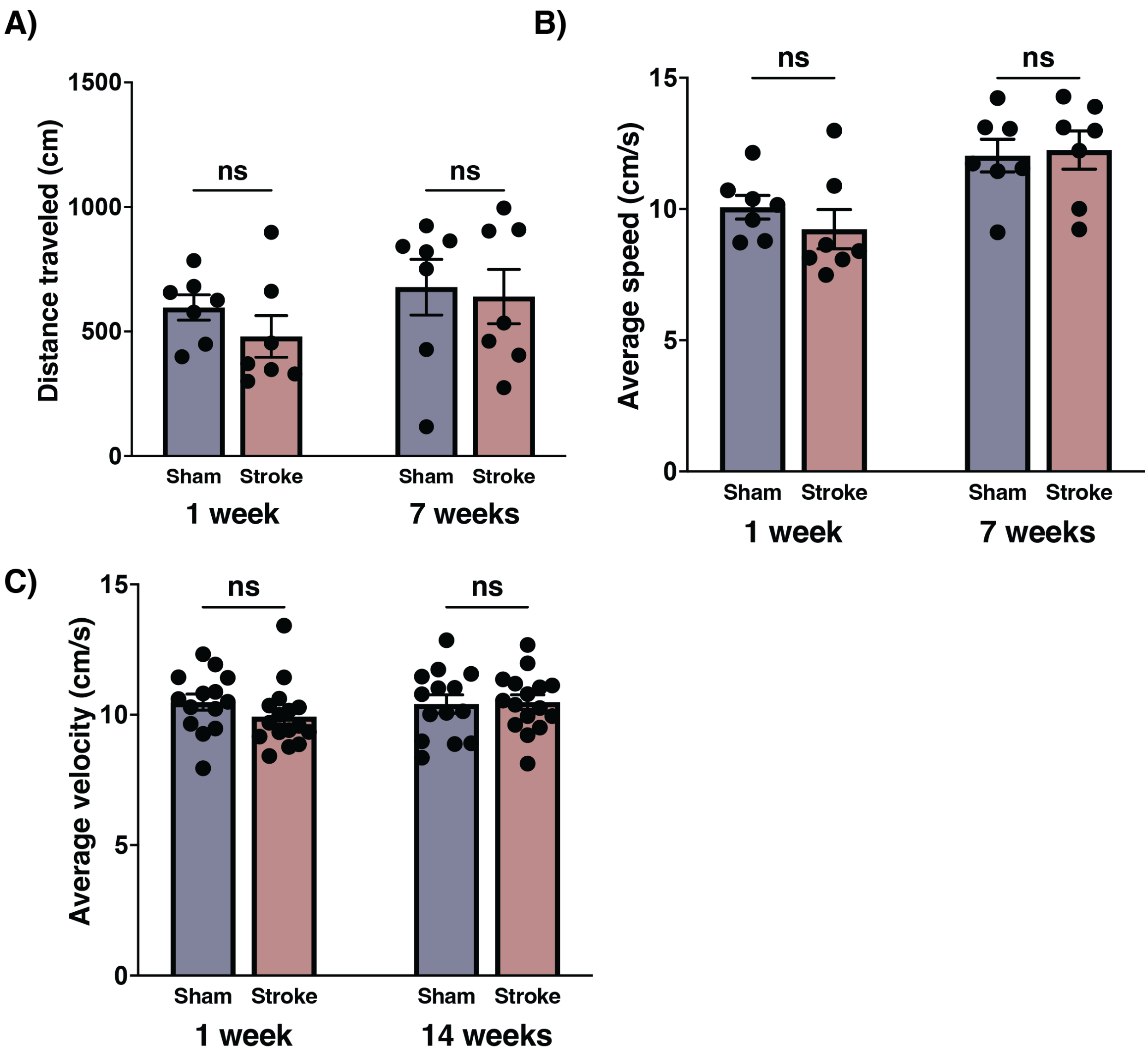


**Supplemental Figure 1: Juvenile DH stroke mice do not have evidence of motor deficit or fatigue**. A) There is no difference in distance traveled on the Barnes Maze spatial paradigm (where DH stroke mice do not develop a deficit) at either 1 or 7 weeks after stroke (n = 7-8/group, Mann-Whitney) B) There is no difference in average speed on the Barnes Maze spatial paradigm at 1 or 7 weeks after stroke (n = 7-8/group, Mann-Whitney). C) There is no difference in average speed on open field testing at 1 or 14 weeks after stroke (n = 14-15/group, Mann-Whitney).

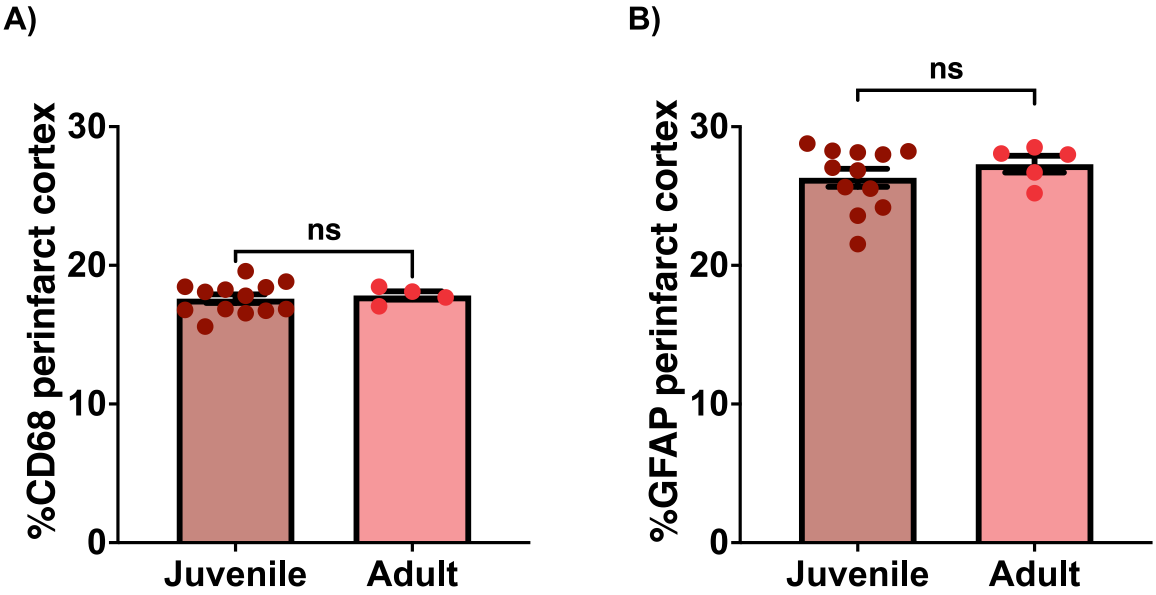


**Supplemental Figure 2: There are no age-related differences in acute innate inflammation in the peri-infarct cortex.** A) At 3 days after stroke, there is no significant difference between juveniles and adults in CD68 staining for activated myeloid cells in the peri-infarct cortex (n = 4 adult, 13 juveniles/group, p = 0.6391, Mann-Whitney). B) At 3 days after stroke, there is no significant difference between juveniles and adults in GFAP staining for reactive astrocyte in the peri-infarct cortex. (n = 4 adult, 13 juveniles/group, p= 0.7214, Mann-Whitney).

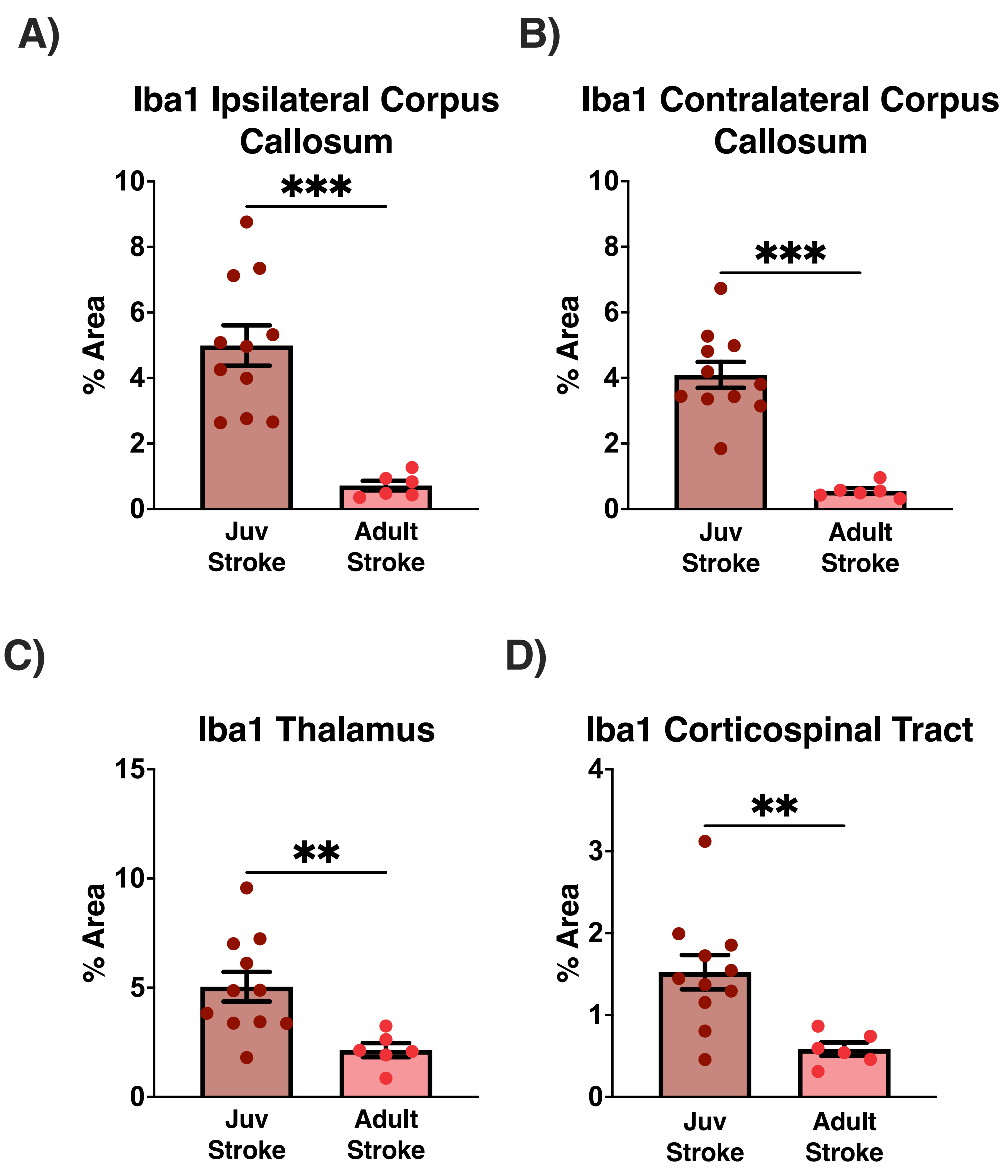


**Supplemental Figure 3: Juvenile mice have significantly greater Iba1s staining at uninjured subcortical gray and white matter regions connected to the infarct.** A) Significantly greater Iba1 staining in the ipsilateral corpus callosum of juvenile stroke mice as compared to adult stroke mice, n = 11 juvenile, 6 adults, p = 0.0002, Mann-Whitney. B) Significantly greater Iba1 staining in the contralateral corpus callosum of juvenile stroke mice as compared to adult stroke mice, n = 11 juvenile, 6 adults, p = 0.0002, Mann-Whitney. C) Significantly greater Iba1 staining in the thalamus of juvenile stroke mice as compared to adult stroke mice, n = 11 juvenile, 6 adults, p = 0.0031, Mann-Whitney. D) Significantly greater Iba1 staining in the corticospinal tract of juvenile stroke mice as compared to adult stroke mice, n = 11 juvenile, 6 adults, p = 0.0048, Mann-Whitney.

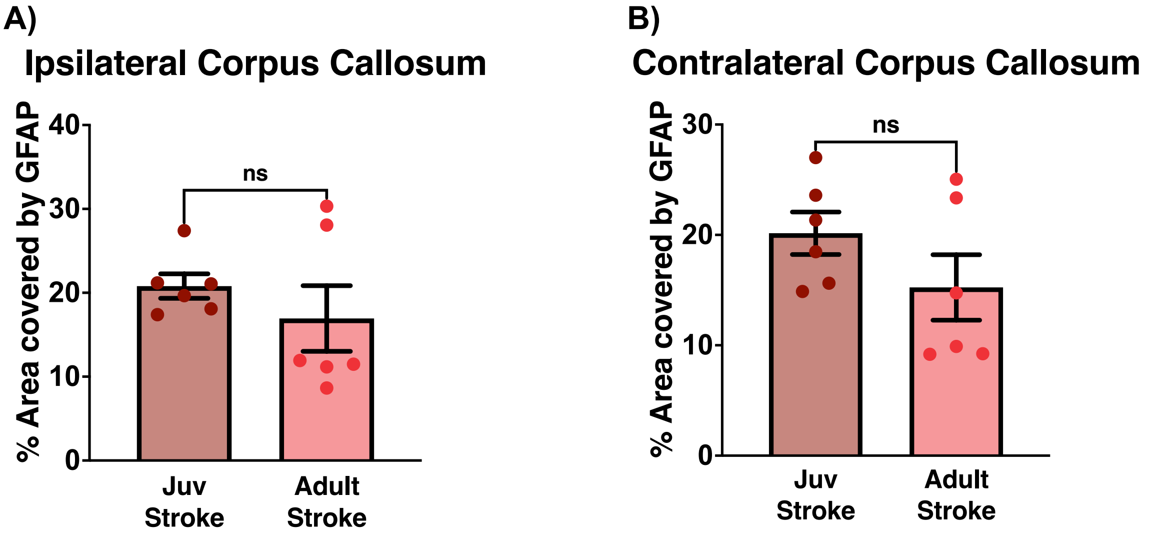


**Supplemental Figure 4: There are no age-related differences in acute astroglial inflammation in the corpus callosum.** A) At 3 days after stroke, there is no significant difference between juveniles and adults in reactive astrocyte staining in the ipsilateral corpus callosum (n = 6/group, p = 0.3939, Mann-Whitney). B) At 3 days after stroke, there is no significant difference between juveniles and adults in reactive astrocyte staining in the contralateral corpus callosum. (n = 6/group, p= 0.1797, Mann-Whitney).
